# Extracellular Vesicle microRNAs From Small Airways Promote Senescence and Fibrosis in COPD

**DOI:** 10.64898/2026.03.30.713627

**Authors:** Justine V Devulder, Peter S Fenwick, Susan Monkley, Lina Odqvist, Louise E Donnelly, Peter J Barnes

**Affiliations:** National Heart and Lung Institute, Imperial College London, Dovehouse Street, SW3 6LY, London, United Kingdom; Translational Sciences & Experimental Medicine, Research & Early Development, Respiratory & Immunology, BioPharmaceuticals R&D, AstraZeneca, Gothenburg, Sweden; Bioscience COPD/IPF, Research and Early Development, Respiratory & Immunology, Biopharmaceuticals R&D, AstraZeneca, Gothenburg, Sweden

**Author notes:** Corresponding author: Justine Devulder. <u>Author contributions:</u> PJB, LED and JVD designed research, JD PSF performed.research, JD and SM analysed the data, LO and SM contributed analytic tools, JD LED PJB and LO wrote the paper.

**Keywords:** Extracellular vesicles, miRNAs, cellular senescence, COPD

## Abstract

**Background:** Chronic obstructive pulmonary disease (COPD) is a chronic lung condition characterised by accelerated lung aging. Extracellular vesicles (EVs), which can be categorised into large EVs (LEVs) and small EVs (SEVs), may play a critical role in intercellular communication. They contribute to the pathogenesis of COPD by transporting and transferring microRNAs (miRNAs). This study profiles cells and EV-associated miRNAs from both healthy and COPD small airway (SA)-epithelial cells and SA-fibroblasts and identifies the biological pathways associated with these miRNAs.

**Methods:** EVs were isolated from conditioned media of healthy and COPD SA-epithelial cells and SA-fibroblasts, both at baseline and following H_2_O_2_ exposure. MiRNAs were extracted from cells and EVs and analysed by small RNA (smRNA) sequencing.

**Results:** SmRNA sequencing of COPD SA-epithelial cells and EVs revealed that four miRNAs were upregulated and fourteen were downregulated in the cells compared to healthy controls. COPD LEVs displayed nine upregulated and ten downregulated miRNAs, while SEVs showed ten upregulated and eleven downregulated miRNAs. Only one miRNA consistently upregulated in COPD SA-epithelial cells, LEVs, and SEVs. The various differentially expressed miRNAs were primarily associated with cellular senescence pathways. In SA-fibroblasts 39 miRNAs were upregulated in COPD compared to healthy cells. 14 miRNAs were upregulated in COPD LEVs and 11 downregulated, whereas SEVs exhibited twenty upregulated and eleven downregulated miRNAs. Overlap was limited, with only three miRNAs consistently upregulated in SA-fibroblasts and EVs. These miRNAs were linked to pathways related to fibrosis and cellular senescence. Furthermore, oxidative stress alters the miRNA profiles detected in cells and EVs differently between cells from healthy individuals and COPD patients.

**Conclusions:** COPD alters miRNA signatures in cells and their EVs, with limited overlap between compartments. These COPD-associated miRNAs are enriched in pathways driving cellular senescence and fibrosis, suggesting a potential role in disease progression.

## Introduction

Chronic obstructive pulmonary disease (COPD) is a progressive inflammatory lung disease characterised by airflow limitation and changes in airway structures leading to chronic bronchitis, small airway disease, and emphysema [1]. It is the third leading cause of death worldwide, with over 390 million people currently living with COPD, posing a significant burden on healthcare systems [2]. Although current therapeutics can help manage disease symptoms, there is no curative treatment for COPD [3].

Premature ageing of the lungs is a key feature of COPD pathophysiology and is associated with increased cellular senescence [4]. Cellular senescence is the cessation of cell division following a defined number of divisions or due to cellular stressors such as oxidative stress. Senescent cells resist apoptosis and undergo various molecular and phenotypical changes, such as increased β-galactosidase activity, enhanced expression of cell cycle inhibitors like p21^CIP1^ (cyclin-dependent kinase inhibitor-1), p16^INK4a^ (cyclin-dependent kinase inhibitor-2A), and p53 (cellular tumor antigen p53), as well as the production of pro-inflammatory factors associated with the senescence-associated secretory phenotype (SASP) [5]. In the lungs of COPD patients, chronic oxidative stress arises from an imbalance between the production of free radicals and a decrease in antioxidant production [6]. Key exogenous sources of oxidative stress include tobacco smoking, occupational dust [7], and indoor biomass fuels [8], which are significant risk factors for the development of COPD. Oxidative stress contributes to the pathogenesis of COPD by stimulating the production of pro-inflammatory factors through the NF-κB pathway, promoting the development of emphysema and fibrosis by reducing antiproteases and producing transforming growth factor (TGF)-β [9]. This chronic oxidative stress leads to the accumulation of senescent cells, including alveolar type (AT)2 cells, endothelial cells, smooth muscle cells, epithelial cells, and fibroblasts [10, 11]. These cells show elevated levels of p21^CIP1^ and p16^INK4a^, along with a decrease in anti-aging molecules such as sirtuins (SIRT) and the production of proinflammatory factors that match the profile of the SASP [12–14]. Additionally, a subset of small airway (SA)-fibroblasts in COPD exhibits a cellular senescent phenotype, which may be associated to mitochondrial dysfunction, an oxidative stress response, and fibrosis [11].

MicroRNAs (miRNAs) are short RNA molecules, typically 19 to 25 nucleotides long, which regulate post-transcriptional silencing of target genes [15]. miRNAs are key regulators of various biological processes and contribute to several diseases, including COPD [16]. Several miRNAs can affect a single gene, and conversely, multiple genes can be impacted by a single miRNA [17]. Profiling studies have revealed differential expression of miRNAs in bronchoalveolar lavage fluid, induced sputum, plasma, and lung tissues from COPD patients compared to healthy non-smokers and smokers [17–20]. Cigarette smoke and oxidative stress alter miRNA expression, potentially affecting cellular senescence and the pathophysiology of COPD. Notably, miR-34a and miR-570 are induced by oxidative stress and COPD SA-epithelial cells display elevated expression compared to cells from healthy donors and smokers [21, 22]. These miRNAs promote cellular senescence in COPD cells by downregulating their target anti-aging molecules, SIRT1 and SIRT6.

Extracellular vesicles (EVs) are particles released from cells that are delimited by a bilayer lipid membrane and cannot replicate independently [23]. They are classified based on their biogenesis and size into large vesicles or microvesicles (LEVs) that originate from the plasma membrane and small vesicles or exosomes (SEVs) that bud off from the endoplasmic reticulum [24]. EVs play a crucial role in intercellular communication as they encapsulate specific cargo, including DNA, RNA, miRNA, or proteins, and transfer it to recipient cells, triggering phenotypic or molecular responses [25]. While LEV and SEV share similarities in their lipid composition, their cargoes are remarkably different due to their distinct biogenesis pathways. Consequently, LEV and SEV can have different effects when internalized in recipient cells [26]. Modifications of EV composition and cargo have been documented in the sputum, bronchoalveolar lavage fluid, and plasma of COPD patients. Recent studies have shown that EVs from SA-epithelial cells in COPD are enriched with miR-34a, which can be transferred to and induce cellular senescence in healthy recipient SA-epithelial cells, highlighting the role of EVs in propagating cellular senescence within the lungs [27]. Oxidative stress can also modify expression of miRNAs within EVs, potentially contributing to cellular senescence and fibrosis in the lungs of individuals with COPD. For instance, bronchial epithelial cells exposed to cigarette smoke produce EVs that are enriched with miR-210, which induces fibroblast differentiation into myofibroblasts [28, 29]. However, there is still limited understanding of the function of miRNAs associated with EVs derived from SA-epithelial cells and SA-fibroblasts and their contributions in the pathophysiology of COPD.

We hypothesise that the miRNAs profiles are dysregulated in SA-epithelial cells and SA-fibroblasts and the EVs produced by them in COPD. We have performed smallRNA (smRNA) sequencing of healthy and COPD SA-epithelial cells and SA-fibroblasts and their respective EVs, at baseline and in response to oxidative stress. We then analysed the pathways associated with these differential miRNAs, focusing particularly on cellular senescence and fibrosis pathways.

## Materials and Methods

### Cell culture

Primary SA-epithelial cells were cultured as monolayers in LHC-9 medium (Invitrogen) on collagen (1% w/v) coated plates. Primary SA-fibroblasts were cultured as monolayers in Dulbecco’s Modified Eagle Medium (DMEM) (Invitrogen) supplemented with 1% v/v penicillin-streptomycin, 1% (w/v) L-glutamine and 1% (w/v) amphotericin B. Cells were extracted from lung tissue from subjects undergoing lung resection surgery at the Royal Brompton Hospital, London. The subjects were matched for age and COPD subjects (Table 1). All subjects gave informed written consent, and the study was approved by the London-Chelsea Research Ethics committee (study 15/SC/0101). Cells were stimulated with 100μM of hydrogen peroxide (Sigma) for 48h.

**Table 1:** Characteristics of study subjects. Patients with COPD were categorised according to Global Initiative for Chronic Obstructive Lung Diseases. Definitions. M, male; F, female; FEV1, forced expiratory volume in 1s; FVC, forced vital capacity; aNumber of cigarettes smoked per day/20 x duration of smoking. Data are expressed as mean ± SEM, *p>0.05. **p>0.01

| Characteristics | Non-COPD SAEC | COPD SAEC | Non-COPD SAF | COPD SAF |
| --- | --- | --- | --- | --- |
| Age (years) | 72.2 ± 2.39 | 74.6 ± 3.48 | 67.2 ± 3.02 | 74 ± 4.25 |
| Sex (M:F) | 02:03 | 04:01 | 02:03 | 04:01 |
| FEV1 (L) | 2.50 ± 0.14 | 1.59 ± 0.13* | 2.55 ± 0.18 | 1.36 ± 0.19** |
| FEV1 (% predicted) | 102.8 ± 5.23 | 57.28 ± 13.59 | 98.8 ± 3.15 | 55.3 ± 17.60 |
| FVC (L) | 3.34 ± 0.22 | 3.36 ± 0.30 | 3.156 ± 0.28 | 3.4375 ± 0.28 |
| FEV1:FVC | 0.752 ± 0.02 | 0.5 ± 0.06** | 0.77 ± 0.04 | 0.47 ± 0.05** |
| Pack-years<sup>a</sup> | 0.0 | 38 ± 11.14** | 0.0 | 42.5 ± 13.15** |

### Isolation of EVs

Isolation of EVs complies with the latest MISEV 2024 guidelines [23]. 2.10^6^ of SA-epithelial cells or SA-fibroblasts were seeded in T75 flasks in 10ml of serum-free cell media depleted of EVs by ultracentrifugation (100000*g* for 2h). After 48h, cell viability was above 95%. Conditioned media was centrifuged at 300g for 10min to eliminate debris, then at 20000g for 30min to isolate large EVs and at 100000g for 2h for small EVs. The resultant pellets were resuspended in phosphate buffered saline (PBS), Qiazol (Qiagen) or RIPA buffer (Sigma) depending on the experiment conducted. The expression of markers of EVs, CD9 (EPR2949, ab92726, Abcam), CD81 (EPR4244, ab232390, Abcam), CD63 (ab68418, Abcam), Alix (ab88388, Abcam, Cambridge UK) and calreticulin (Biotechne, MAB38981, Minneapolis USA) were analysed by Western Blot. EV concentration was measured by NanoFCM (NanoFCM, Xiamen, China).

### Characterisation of EVs using nanoflow cytometry analysis

Large (LEVs) and small EVs (SEVs) were analysed by NanoFCM (NanoFCM, Xiamen, China) for particle concentrations and size distribution. The instrument was calibrated for concentrations using polystyrene beads and for size distribution using Silica Nanosphere cocktail (NanoFCM Inc., S16M-Exo). Any particles that passed by the detector during a 1 min interval were recorded in each test. All samples were diluted in 0.22μm-filtered PBS to attain a particle count within the optimal range of 2000-12000/min. Using the calibration curve, EVs concentration and size were determined on the NanoFCM software (NanoFCM Profession V1.0).

### RNA isolation

Cells, LEVs and SEVs were resuspended in Qiazol® (Qiagen). Total RNA, including small RNAs, were extracted using the miRNeasy kit (Qiagen) according to the manufacturer’s instructions. RNAs was quantified by NanoDrop (Thermofisher Scientific, Waltham, USA). 100ng of total EVs RNA and 500ng of cells RNA was reverse transcribed using the Taqman total RNA including small RNA reverse Transcription kit (Applied Biosystems, Waltham, Massachusetts, USA) or TaqMan™ Advanced miRNA cDNA Synthesis Kit (Applied Biosystems, Waltham, Massachusetts, USA). miRNA levels were detected by TaqMan MicroRNA Assay (hsa-miR-376a-3p, hsa-miR-376a-5p, hsa-miR-204-5p, hsa-miR-137-3p) (Life Technologies, Waltham, Massachusetts, USA). RNU48 was detected as the endogenous control for miRNA detection in cells and in EVs as its expression is not modified by any stimulation used in our study. After the reactions, the CT values were determined using fixed-threshold settings. The relative fold difference was calculated using the _2-ΔΔCt._

### smRNA-sequencing and statistical analysis

RNA isolated from cells and their derived EVs was quantified using Qubit RNA Assay Kits, and its integrity was measured by an Agilent Bioanalyzer (carried out by Source BioScience, Cambridge, UK). Libraries were prepared using the NEBNext® Ultra^TM^ II Directional RNA Library Prep Kit according to the manufacturer’s protocol. During this process, the libraries were indexed using NEBNext® Multiplex Oligos for Illumina® (Source BioSciences). The prepared libraries were quantified via a fluorometric method involving a Promega QuantiFluor dsDNA assay; and qualified using electrophoretic separation on the Agilent TapeStation 4200. All samples were sequenced on the NovaSeq and data was returned as raw fastQ data. Data received was run through a pre-defined smRNA-seq pipeline which includes quality control (QC), adapter sequence trimming of raw reads by Atropos, STAR to align against the genome (hg38) and other smRNA detection by SeqCluster. miRNAs were detected using miraligner tool with miRbase (21 release, http://www.mirbase.org/) as the reference miRNA database, the quantification of known smRNA was carried out by SeqBuster. Count matrices were generated for miRNA and isomiRs by combining data for each sample with miRNAs as rows and samples as column. FastQC was used for QC and multiqc for reporting. The miRNA count matrix was used for differential expression analysis in R using DESeq2 package. Results were later filtered for adjusted p-values (p.adj) threshold (or false discovery rate, FDR) of <0.1.

Qiagen Ingenuity Pathway Analysis (IPA) software, Pubmed and miRDB databases were used to investigate differentially expressed miRNA datasets and mRNA targets. miRNA-mRNA relationships and gene predictions were prioritised based on cellular senescence and fibrosis pathways links before visualisation. IPA’s microRNA Target Filter uses content from Tarbase, miRecords, TargetScan and published literature while DIANA-miRpath KEGG and GO analysis takes place using Tarbase.

Data are expressed as mean ± SEM. Results were analysed using Mann-Whitney. GraphPad Prism 9.2 software (GraphPad software, La Jolla, CA) was used for analysis. Values of p≤0.05 were considered statistically significant.

## Results

### COPD small airway epithelial cells, but not fibroblasts, release higher amount of EVs than healthy cells

EVs derived from SA-epithelial cells and SA-fibroblasts from healthy donors and COPD patients were characterized using Western blot, NanoFlow cytometry, and transmission electron microscopy (TEM) (**Figure 1**). Canonical EV markers including CD9, CD81, CD63, and Alix were detected in all EVs. EpCam was specific to SA-epithelial cells derived EVs, while CD90 was exclusive to SA-fibroblasts-derived EVs. The absence of calreticulin indicated purity of the preparations (**Figure 1A**). NanoFCM analysis revealed average sizes of 89.5 ± 2.4 nm for SA-epithelial cells LEVs and 73.7 ± 0.9 nm for SEVs, compared to 84.8 ± 2.2 nm and 78.1 ± 1.5 nm for SA-fibroblasts LEVs and SEVs, respectively (**Figure 1B**). No significant differences in size distribution were observed between healthy and COPD donors or between cell types. TEM confirmed the double-membrane morphology and cup-shaped structure of the vesicles (**Figure 1C**). Quantification demonstrated that COPD SA-epithelial cells produced significantly more EVs than healthy cells. COPD SA-fibroblasts also seems to release more LEVs than healthy cells, although SEV levels remained similar (**Figure 1D**).

**Figure 1:**
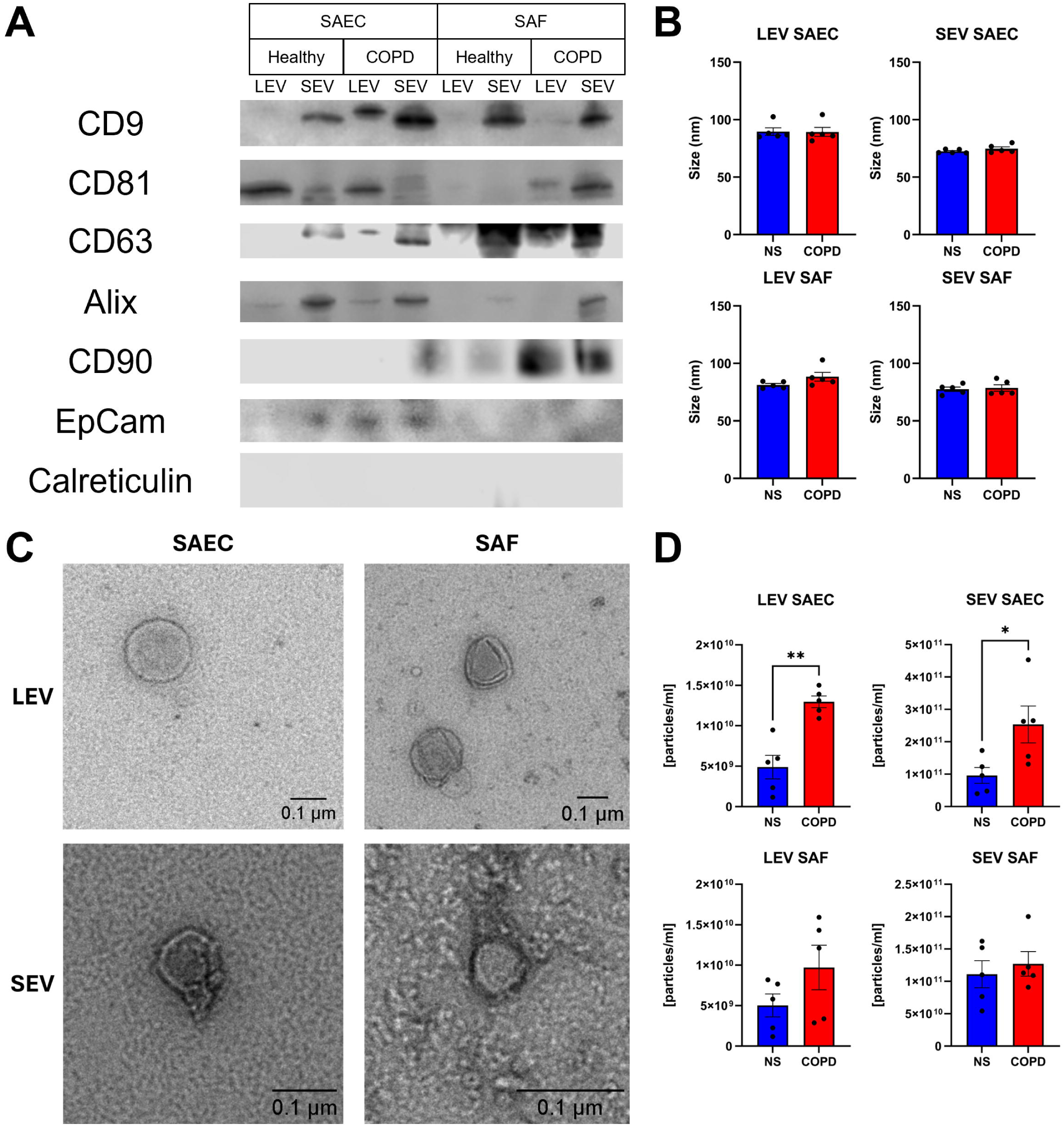
Characterisation and quantification of EVs produced by healthy and COPD SA-epithelial cells and SA-fibroblasts. Large and small EVs were isolated from 48 hours-conditioned cell media from healthy or COPD SA-epithelial cells or SA-fibroblasts. **A.** Markers of EVs, of epithelial cells and fibroblasts were measured by Western Blot. **B.** The size of LEVs and SEVs was analysed by NanoFCM. **C.** The shape and size of EVs were confirmed by TEM. **D.** The concentration of EVs were analysed by NanoFCM. Each point represents a different subject. Data are mean ± SEM, analysed by Mann-Whitney test (n=5), *p<0.05, **p<0.01

### SA-epithelial cells, SA-fibroblasts, and Derived EVs Exhibit Distinct miRNA Signatures in COPD

To investigate whether miRNAs contained in EVs derived from SA-epithelial cells and SA-fibroblasts differed between COPD and controls, we performed small RNA sequencing on both parental cells and their derived EVs (**Figure 2 and 3**). In SA-epithelial cells, 40.7% of smRNAs were miRNAs, compared to 15–18% in EVs, with rRNAs enriched in EVs (**Figure 2A**). In total, 2971 miRNAs were detected in SA-epithelial cells, 1050 in LEVs and 953 in SEVs (**Figure 2B**). The analysis of miRNA profiles revealed a distinct separation between the profiles of EVs and parental SA-epithelial cells **(Figure 2C**). In COPD SA-epithelial cells, four miRNAs were upregulated and fourteen downregulated compared to healthy controls. COPD LEVs exhibited nine upregulated and ten downregulated miRNAs, while SEVs showed ten upregulated and eleven downregulated (**Figure 2D, 2E and 2F, Table S1**). Overlap between dysregulated miRNAs in cells and EVs was minimal, with only miR-376a-3p consistently upregulated in COPD SA-epithelial cells, LEVs, and SEVs (**Figure 2G**). To validate these results, we measured the expression of miR-376a-3p using RT-qPCR in healthy and COPD SA-epithelial cells, LEVs, and SEVs. The results from the smRNA sequencing were confirmed, as miR-376a-3p expression was significantly increased in COPD cells and LEVs, with a non-significant increase observed in COPD SEVs (**Figure 2H**).

**Figure 2:**
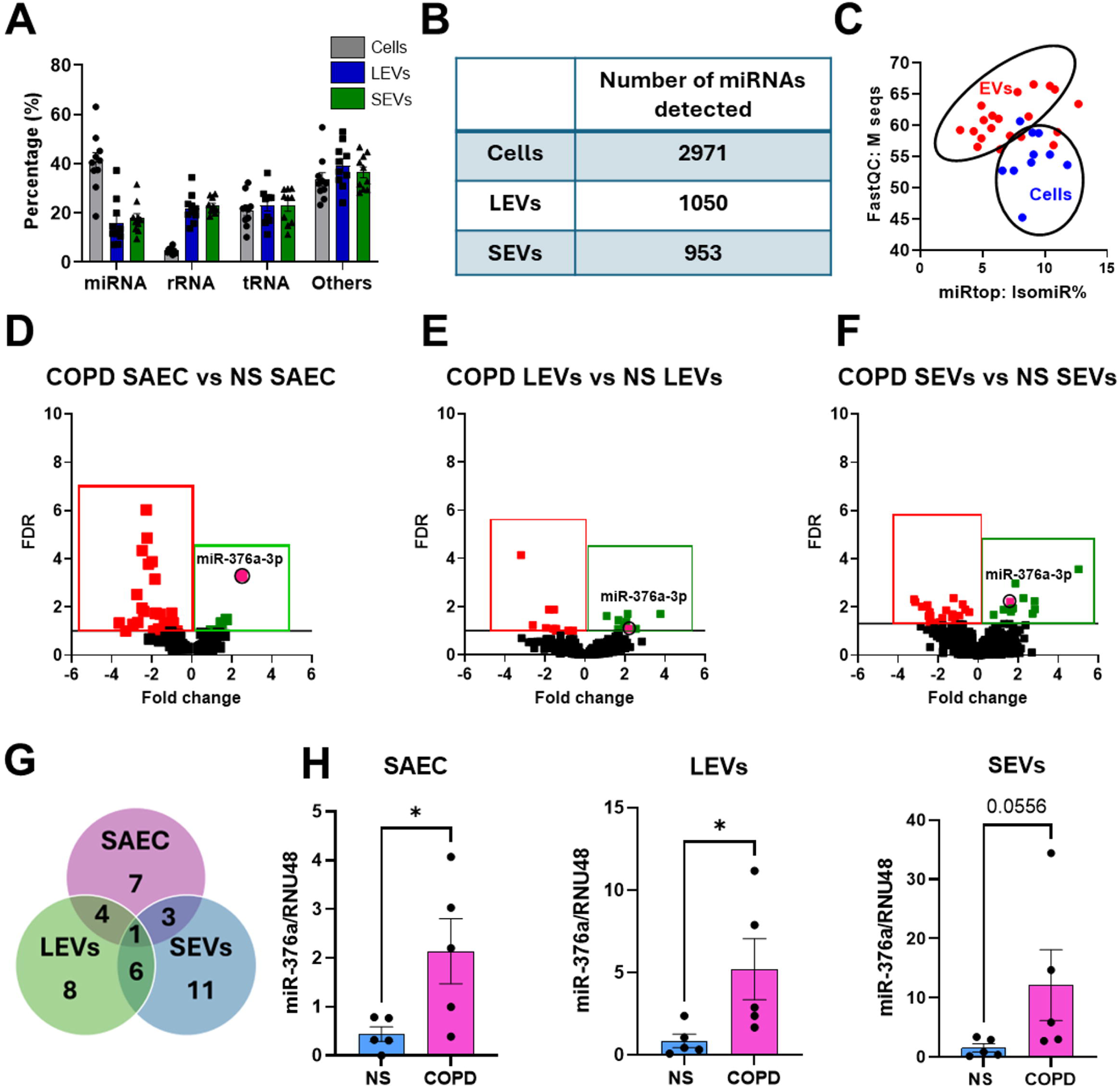
Dysregulation of miRNA Expression in COPD SA-epithelial cells and associated EVs. Small RNA sequencing was conducted on healthy and COPD SA-epithelial cells (SAEC) and their derived large and small EVs. **A.** The proportions of miRNA, rRNA, tRNA and other types of smRNA were characterised and **B.** the numbers of miRNAs detected in both SAEC and EVs were assessed. **C.** The percentage of IsomiR (variants of miRNAs) and Mseq (sequence count) were plotted to examine the miRNAs profiles in cells and EVs. **D-F.** Volcano plots were used to compare miRNA expression between healthy and COPD cells and EVs associated. **G.** A Venn-diagram illustrated the overlap of dysregulation miRNA between healthy and COPD cells and EVs. **H.** The sequencing data were validated by RT-qPCR for miR-376a-3p (n=5). Padj value<0.1. Data are represented as mean ± SEM and analysed by Mann-Whitney test. *p<0.05

**Figure 3:**
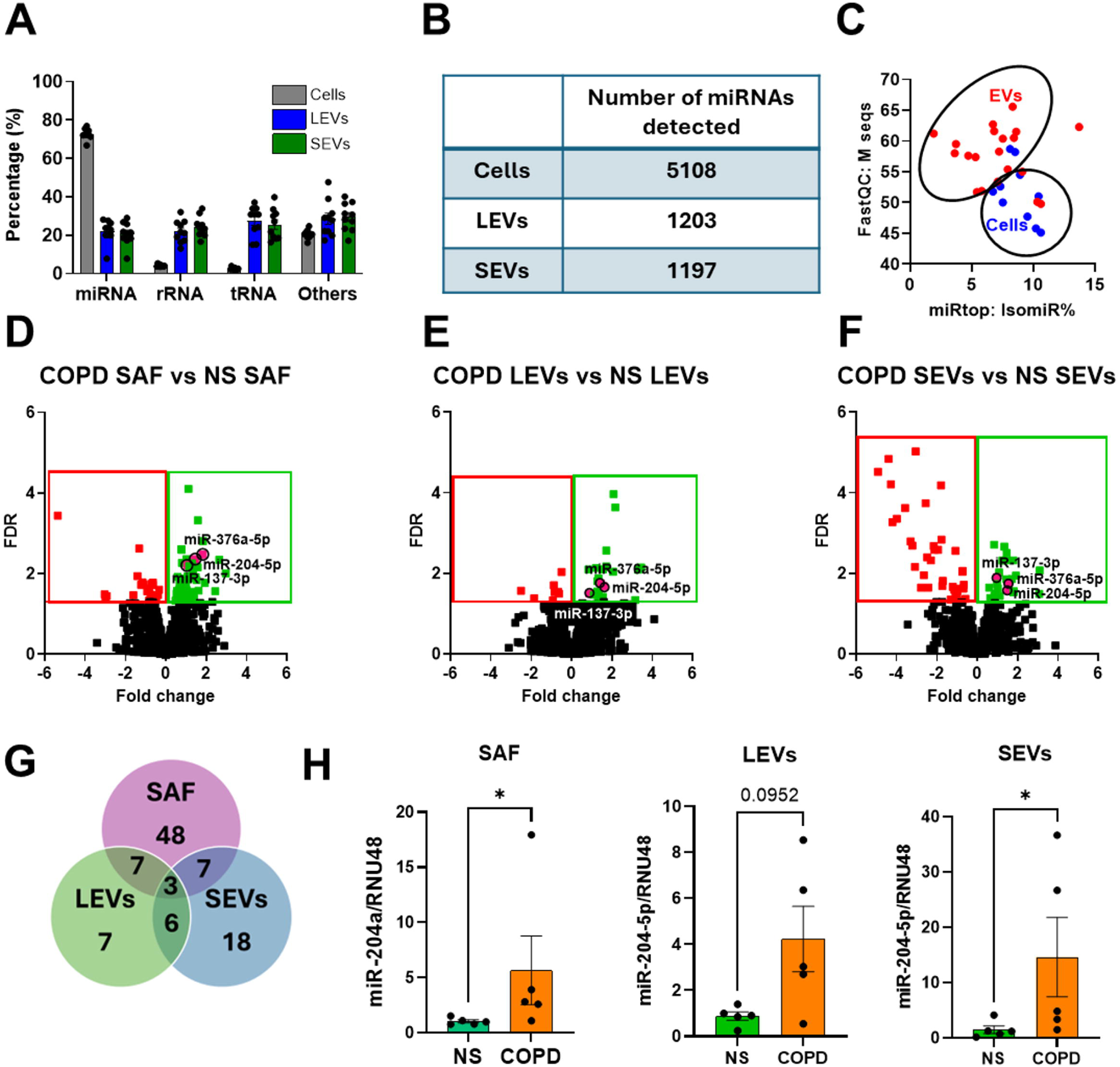
Dysregulation of miRNA Expression in COPD SAF and Associated EVs. Small RNA sequencing was conducted on healthy and COPD SAF and their derived large and small EVs. **A.** The proportions of miRNA, rRNA, tRNA and other types of smRNA were characterised and **B.** the numbers of miRNAs detected in both SAF and EVs were analysed. **C.** The percentage of IsomiR (variants of miRNAs) and Mseq (sequence count) were plotted to examine the miRNAs profiles in cells and EVs. **D-F.** Volcano plots were used to compare miRNA expression between healthy and COPD cells and EVs associated. **G.** A Venn-diagram illustrated the overlap of dysregulation miRNA between healthy and COPD cells and EVs. **H.** The sequencing data were validated by RT-qPCR for miR-204a-5p (n=5). Padj value<0.1. Data are represented as mean ± SEM and analysed by Mann-Whitney test. *p<0.05

In SA-fibroblasts, 72.7% of smRNAs were miRNAs, compared to approximately 22.06 ± 1.8% in LEVs and 20.7 ± 1.8% in SEVs (**Figure 3A**). SA-fibroblasts contained 5108 miRNAs, while LEVs and SEVs contained 1203 and 1197, respectively (**Figure 3B and 3C**). COPD SA-fibroblasts exhibited thirty-nine upregulated and thirteen downregulated miRNAs (**Figure 3D**). COPD LEVs had fourteen upregulated and six downregulated, while SEVs had twenty upregulated and eleven downregulated miRNAs (**Figure 3E and 3F, Table S2**). Overlap was limited, with only three miRNAs (miR-376a-5p, miR-204-5p, and miR-137-3p) consistently upregulated in SA-fibroblasts and EVs (**Figure 3G**). These results were validated by RT-qPCR, showing that miR-204-5p was significantly upregulated in COPD SA-fibroblasts and SEVs, with a non-significant increase in LEVs (**Figure 3H**). Expression of miR-376a-5p was increased in COPD SA-fibroblasts and SEVs, but it was not detected in healthy or COPD LEVs (**Figure S1A**). Expression of miR-137-3p seemed to increase in COPD SA-fibroblasts; however, it was undetected in LEVs and SEVs (**Figure S1B**). Overall, these findings confirm that the miRNA profiles differ according to disease status and are cell specific. Additionally, we demonstrated that the miRNA profiles detected in COPD cells differ from those observed in EVs, suggesting that the packaging of miRNAs into EVs is an active process rather than a passive one. This may account for the differing miRNA profiles observed in cells compared to EVs.

### Dysregulated miRNAs in COPD cells and EVs are involved in cellular senescence and fibrosis pathways

We then analysed the genes and pathways associated with the significantly dysregulated miRNAs using Ingenuity Pathway analysis (KEGG and Gene Ontology). In SA-epithelial cells and their EVs, dysregulated miRNAs were associated with phosphatase and tensin homolog (PTEN) signalling, p53 signalling, Signal transducer and activator of transcription (STAT)3 signalling, phosphatidylinositol 3-kinase (PI3K)/ protein kinase B (AKT) signalling, autophagy, cell cycle regulation, and SASP, such as CXCL8 signalling **(Figures 4A, 4B, and 4C**). Notably, miR-136-5p was found to be upregulated in COPD SA-epithelial cells and LEVs, where it plays a role in cellular senescence by targeting Pyruvate Dehydrogenase Kinase (PDK)4 (**Figure 4D**) [30]. The dysregulated miRNAs in COPD-derived SA-fibroblasts and their associated EVs were found to be involved not only in senescence pathways, such as cell cycle regulation, p53 and PTEN signalling, and PI3K/AKT signalling, but also in fibrosis-associated pathways. These include the pulmonary fibrosis signalling pathway, wound healing pathways, epithelial-mesenchymal transition (EMT) regulation, focal adhesion kinase (FAK) signalling, and epithelial adherens junction signalling (**Figures 5A, 5B, and 5C**). miR-137-3p promote senescence via CDK6 and fibrosis via COL5A1 while miR-204-5p promotes senescence by targeting CDC25 and fibrosis through FBN2, FRS2 and TGFβR2 [31, 32] (**Figure 5D**).

**Figure 4:**
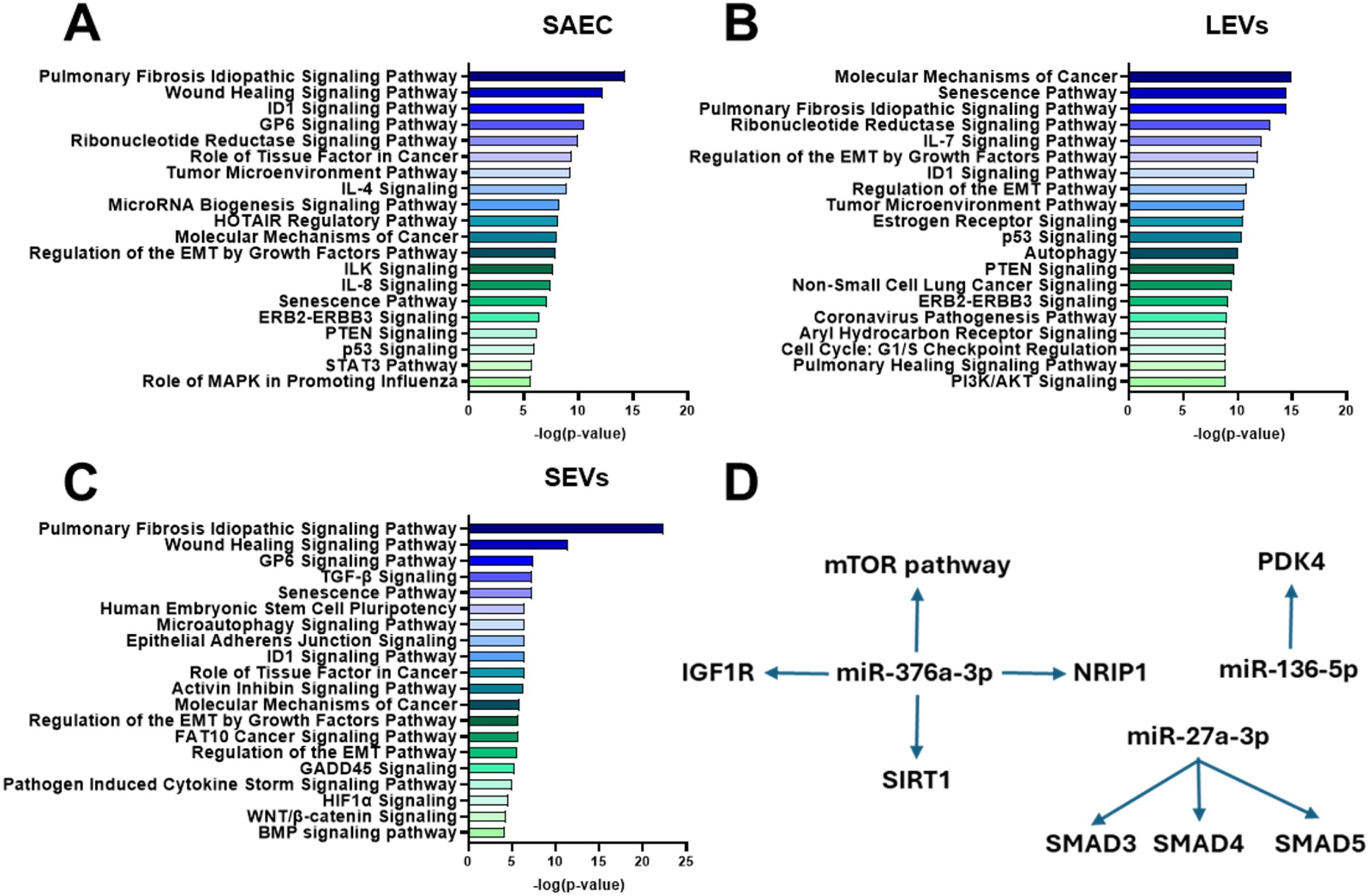
miRNA dysregulated in COPD SAEC and EVs are involved in cellular senescence pathways. The targets and pathways controlled by miRNAs dysregulated in COPD compared to healthy were analysed by Ingenuity Pathway analysis software (KEGG and GO pathway analysis) in **A.** SAEC, **B.** LEVs and **C.** SEVs. **D.** The targets of significantly differentially expressed miRNAs (miR-376a-3p, miR-136-5p, and miR-27a-3p) were identified within the context of cellular senescence pathways.

**Figure 5:**
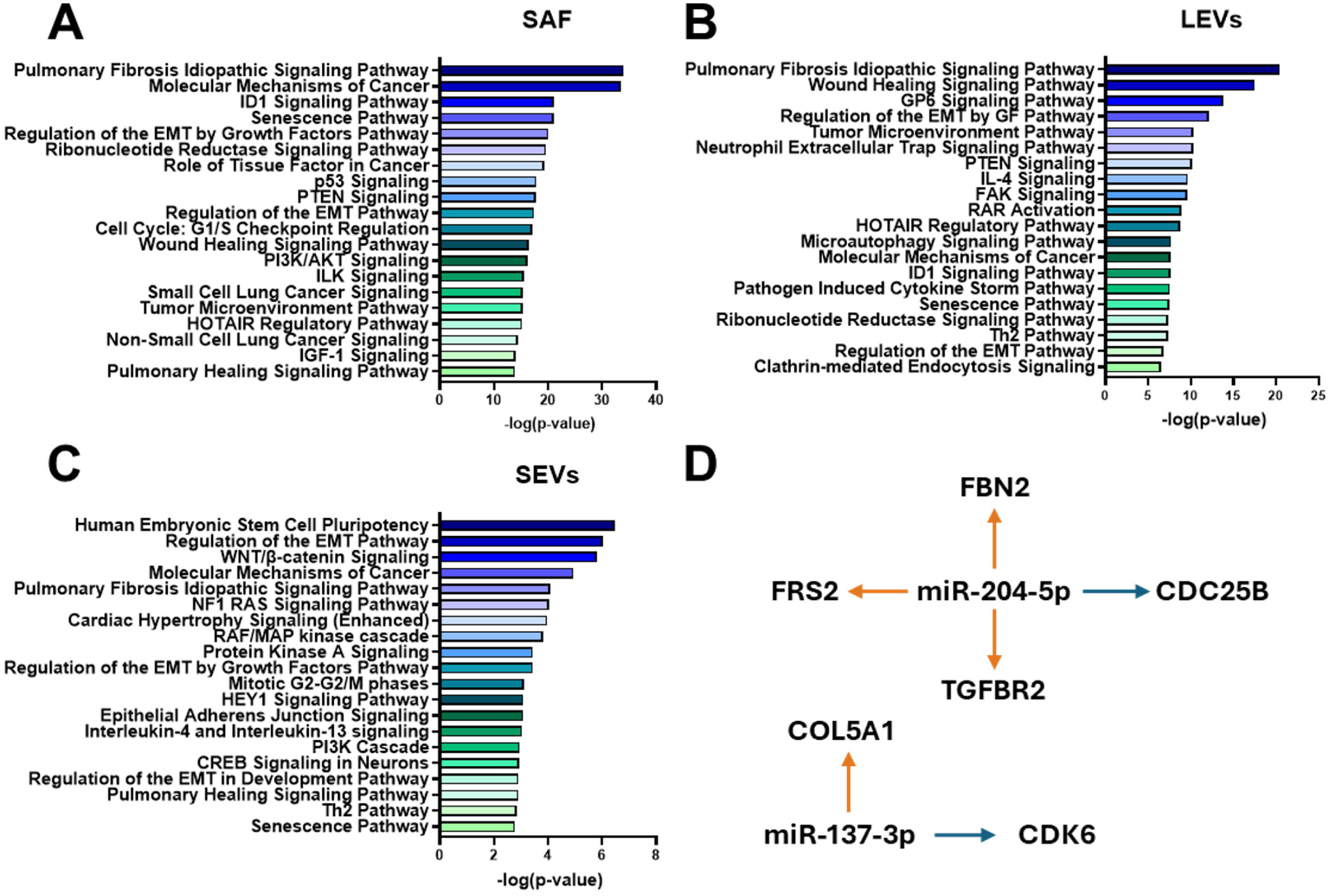
miRNA dysregulated in COPD SAF and EVs are involved in cellular senescence pathways. The targets and pathways controlled by miRNAs dysregulated in COPD compared to healthy were analysed by Ingenuity Pathway analysis software (KEGG and GO pathway analysis) in **A.** SAF, **B.** LEVs and **C.** SEVs. **D.** The targets of significantly differentially expressed miRNAs (miR-137-3p, miR-204-5p) were identified within the context of cellular senescence pathways.

Overall, these findings suggest that dysregulated miRNAs in cells and EVs contribute to the induction of cellular senescence and fibrosis, potentially participating in the pathophysiology of COPD.

### Oxidative stress modulates miRNA Expression in COPD SA-epithelial cells and SA-fibroblasts and their packaging into EVs

To assess the impact of oxidative stress on the expression of miRNA, SA-epithelial cells and SA-fibroblasts were cultured with hydrogen peroxide (H_2_O_2_) for 48h and smRNA sequencing was performed on cells and derived-EVs. In SA-epithelial cells, proportions of smRNA species were similar to baseline (**Figure S2A**). A total of 2927 miRNAs were detected in SA-epithelial cells, 734 in LEVs and 1101 in SEVs (**Figure S2B**). Profiles remained distinct between cells and EVs (**Figure S2C**). Under oxidative stress, COPD SA-epithelial cells showed two downregulated and one upregulated miRNA (**Figure S2D**); COPD LEVs had three upregulated and eight downregulated (**Figure S2E**); SEVs had ten upregulated and sixteen downregulated (**Figure S2F**). Overlap remained minimal, with miR-376a-3p consistently upregulated in COPD SA-epithelial cells and EVs under both baseline and oxidative stress conditions when compared to healthy cells and EVs (**Figure S2G**). In SA-fibroblasts, similar trends were observed, with two miRNAs, miR-22-5p and miR-210-3p, consistently upregulated in COPD SA-fibroblasts and EVs under oxidative stress (**Figure S3**).

To further investigate the effects of H_2_O_2_ on miRNA expression and packaging, we conducted a comparative analysis between baseline and oxidative stress conditions for both healthy and COPD cells and their EVs (**Table S3 and Table S4**). Comparisons revealed oxidative stress-induced-changes differed between healthy and COPD cells (**Figure 6, Table S3 and S4**). miR-4448 and miR-3178 were downregulated in COPD SA-epithelial cells compared with healthy cells at baseline, but their expression was unaffected by H₂O₂ treatment in COPD cells. This reflects a selective downregulation of these miRNAs in healthy SA-epithelial cells following oxidative stress. Additionally, miR-12135 was downregulated in COPD SA-epithelial cells only at baseline; however, H₂O₂ treatment increased its expression in COPD cells, an effect not observed in healthy cells (**Figure 6A**). EVs exhibited distinct stress-dependent responses. miR-181c-3p was increased in COPD LEVs compared with healthy LEVs following H₂O₂ treatment, but not at baseline, due to reduced expression in healthy LEVs under oxidative stress. H_2_O_2_ also impacted the expression of miRNAs on SA-fibroblasts and associated EVs (**Figure 6B**). Some miRNAs, such as miR-204-5p, were consistently upregulated in COPD SA-fibroblasts and derived EVs at baseline and under oxidative stress, whereas others, including miR-17-5p, were downregulated in COPD cells only at baseline. This was driven by oxidative stress-induced downregulation in healthy, but not COPD, fibroblasts (**Figure 6B**).

Overall, miRNA dysregulation in COPD reflects both disease-specific and oxidative stress-dependent effects, which may influence EV miRNA packaging and contribute to COPD pathophysiology.

## Discussion

Cellular senescence and fibrosis are central to COPD and drive small airway disease. EVs are increasingly implicated in COPD pathophysiology, including the induction of paracrine senescence in small airway (SA) epithelial cells [33]. Here, we show that EVs derived from SA-epithelial cells and SA-fibroblasts from COPD patients carry distinct miRNA signatures enriched for pathways linked to senescence and fibrosis. These findings emphasise the contribution of EV-associated miRNAs to COPD and highlight their potential as biomarkers and therapeutic targets.

COPD is linked to dysregulation of miRNAs contained in EVs. Elevated EV numbers have been reported in plasma, bronchoalveolar lavage (BAL), and sputum of COPD patients versus healthy individuals, correlating with systemic inflammation, exacerbation risk, and pulmonary function [34–37]. Altered miRNA expression in cells and EVs has been described in BAL fluid (BALF), sputum, and plasma, though results vary across studies [38]. For example, COPD BALF-EVs have been reported to contain higher miR-122-5p, miR-449c-5p, miR-10a-5p, and miR-155-5p than healthy EVs [39], yet miR-122-5p has also been found decreased in COPD EVs from BALF, lung tissue, sputum, and blood [40, 41]. Similarly, EVs isolated from different sources, such as BALF and plasma, exhibit different miRNA profiles These studies highlight the need for a deeper understanding of the cellular mechanisms that alter EV production and content in COPD. We therefore profiled miRNAs in EVs from SA-epithelial cells and SA-fibroblasts.

Our data indicate that COPD SA-epithelial cells and SA-fibroblasts, and their EVs, exhibit modifications in their miRNA cargo compared with healthy controls. miR-376a-3p, upregulated across COPD SA-epithelial cells and corresponding EVs, has been linked to ageing phenotypes such as Hutchinson–Gilford progeria; its inhibition restores proliferative capacity and reduces senescence markers in progeria fibroblasts [42]. Although direct targets remain to be validated, predictions implicate IGF1R, NRIP1, the mTOR pathway, and SIRT1. miR-204-5p, consistently elevated in COPD SA-fibroblasts and their EVs, has roles in senescence, via CDC25B, and fibrosis through TGFβ signalling and regulation of fibrillin and fibronectin [43]. The link between fibrosis and senescence in COPD is supported by Wrench *et al*., who identified senescent fibroblasts from COPD lungs expressing genes involved in mitochondrial dysfunction and profibrotic pathways [11]. Our previous work showed that miR-34a, but not miR-570, was upregulated in COPD SA-epithelial cells which was involved in the induction of cellular senescence when transferred into healthy recipient SA-epithelial cells [33]. In agreement with this, miR-34a was detected in SA-epithelial cells and SA-fibroblasts and in their EVs and average copies of miR-34a detected was higher in COPD than healthy donors but it did not reach statistical significance, likely reflecting differences in EV isolation conditions and detection sensitivity across studies. Using a highly specific single-cell microfluidic platform, Ho *et al*. recently confirmed miR-34a upregulation in both COPD SA-epithelial cells and SA-fibroblasts, aligning with the trend observed here [44]. Together, our findings support the dysfunction of SA-epithelial cells and fibroblasts in COPD and suggest that miRNAs associated with EVs can simultaneously induce cellular senescence and fibrosis, contributing to the development and progression of small airway disease [45].

EVs mediate intercellular communication by delivering active biomolecules to recipient cells, leading to phenotypic and molecular responses [25]. miRNAs within EVs act as shuttles for genetic exchange [46]. Analysis by microarray techniques has shown differences in miRNA abundance between cells and EVs. However, the analysis of these miRNAs by RT-qPCR suggested that the higher the expression of specific microRNAs was in cells, the higher its expression was in EVs [47, 48] suggesting that miRNA expression in EVs reflects that in cells. However, sequencing studies ranking miRNAs by read counts reveal that while a few miRNAs are highly ranked in both cells and EVs, the majority are predominantly expressed either in cells or in EVs, indicating non-random active sorting [49]. Moreover, distinct miRNA profiles have been identified between healthy and COPD LEVs and SEVs, indicating different sorting mechanisms. Multiple sorting mechanisms have been proposed, including RNA-binding proteins (e.g., Major Vault Protein, Y-Box binding proteins), components of the miRNA machinery (e.g., Argonaute-2), membrane regulators (e.g., caveolin-1, neutral sphingomyelinase-2), and EV biogenesis proteins (e.g., Alix) [50–53]. Our comparisons of miRNA profiles between healthy and COPD cells and their EVs support active sorting, as comparisons between the miRNA profiles of healthy and COPD EVs and cells have demonstrated distinct profiles. Defining how sorting specificity is altered in COPD may explain aberrant EV packaging and would be valuable for future research.

Oxidative stress, a hallmark of COPD, arises from excess production of reactive species and impaired antioxidant defences [6]. Oxidative stress is elevated in COPD lungs and in current and former smokers and persists despite smoking cessation [54]. This increased oxidative stress plays a significant role in triggering cellular senescence by modifying the expression of microRNAs (miRNAs): in both healthy and COPD cells, it induces miR-34a via PI3K activation and PTEN downregulation [22]. Cigarette smoke also increases miR-21 in bronchial epithelial cells, suppressing PTEN and activating Akt [55]. Oxidative stressors (e.g., hydrogen peroxide, ethanol, sulforaphane) increase EV release and alters the RNA profiles associated with these vesicles [56–58]. Next-generation sequencing of EVs from smoke-exposed SA-epithelial cells identified the upregulation of five miRNAs and the downregulation of three miRNAs [59]. Moreover, oxidative stress can promote miRNA loading into EVs by modulating interactions with proteins such as caveolin-1 and hnRNPA2B1 [60]. Consistent with this, we find that hydrogen peroxide alters miRNA expression in both cells and EVs, with distinct patterns in healthy donors versus COPD patients. Prior work shows that Nrf2-mediated antioxidant responses are activated in healthy and smoker cells but are diminished in COPD, exacerbating lung damage [61]. However, the specific impact of oxidative stress on miRNA expression across cells and EVs in COPD had not been defined. To our knowledge, our study is the first to demonstrate disease-specific and compartment-specific effects of oxidative stress on miRNA profiles in paired cells and EVs from SA-epithelial cells and SA-fibroblasts.

This study has limitations. We detected diverse small RNAs in EVs from healthy and COPD SA-epithelial cells and SA-fibroblasts, but many small RNAs, including miRNAs, remain unidentified. Comprehensive EV profiling, including proteins, lipids, and metabolites, will be important, as non-RNA cargo may also contribute to COPD pathophysiology. The miRNAs identified in our analysis are often linked to multiple biological pathways and could play a role in COPD through these interacting pathways. Some of these miRNAs have predicted targets associated with senescence pathways, while others are linked to fibrosis pathways. Future research should focus on validating the targets of these miRNAs that are connected to senescence and fibrosis pathways to better understand their specific mechanisms of action. Finally, functional transfer studies are required to establish how EV-associated miRNAs propagate senescence and fibrosis in recipient SA-epithelial cells and fibroblasts.

In conclusion, EVs (LEVs and SEVs) from SA-epithelial cells and SA-fibroblasts exhibit distinct, disease-modified miRNA signatures in COPD. These miRNAs converge on pathways regulating cellular senescence and fibrosis, offering mechanistic insight into small airway disease. Oxidative stress further reshapes miRNA expression and EV cargo in a disease-dependent manner, suggesting a maladaptive stress response that may amplify lung injury. Together, our data position EV-associated miRNAs as promising biomarkers and therapeutic targets for intervening on cellular senescence and fibrosis in COPD.

