## Supplemental Figures for "Extracellular Vesicle microRNAs From Small Airways Promote Senescence and Fibrosis in COPD"

**Supplementary materials for**

### Altered miRNAs signatures in small airway cells and extracellular vesicles drive cellular senescence and fibrosis in COPD

Justine V Devulder[1], Peter S Fenwick[1], Susan Monkley[2], Lina Odqvist[3], Louise E Donnelly[1], Peter J Barnes[1]

[1] National Heart and Lung Institute, Imperial College London, Dovehouse Street, SW3 6LY, London, United Kingdom

[2] Translational Sciences & Experimental Medicine, Research & Early Development, Respiratory & Immunology, BioPharmaceuticals R&D, AstraZeneca, Gothenburg, Sweden

[3] Bioscience COPD/IPF, Research and Early Development, Respiratory & Immunology, Biopharmaceuticals R&D, AstraZeneca, Gothenburg, Sweden

| SAEC | | | | | |
|---|---|---|---|---|---|
| Upregulated miRNA | | | Downregulated miRNA | | |
| Name | Log2Change | padj | Name | Log2Change | padj |
| hsa-miR-376a-3p | 2.48 | 5.08E-04 | hsa-let-7a-5p | -0.75 | 1.00E-01 |
| hsa-miR-323a-3p;iso_5p:G | 1.72 | 3.34E-02 | hsa-let-7c-5p | -0.92 | 4.71E-02 |
| hsa-miR-136-5p | 1.47 | 4.71E-02 | hsa-let-7i-5p | -1.01 | 1.82E-02 |
| hsa-miR-29a-5p | 0.98 | 1.00E-01 | hsa-miR-218-5p | -1.12 | 1.00E-01 |
| | | | hsa-miR-143-3p | -2.28 | 1.82E-02 |
| | | | hsa-miR-12136 | -1.83 | 1.86E-02 |
| | | | hsa-miR-10a-5p;iso_5p:t | -1.85 | 7.12E-04 |
| | | | hsa-miR-3178;iso_snp:1AG;iso_5p:GC | -1.91 | 1.00E-01 |
| | | | hsa-miR-10a-5p | -1.99 | 1.34E-04 |
| | | | hsa-miR-214-3p | -2.49 | 1.13E-02 |
| | | | hsa-miR-183-5p;iso_5p:t | -1.38 | 4.71E-02 |
| | | | hsa-miR-3182 | -1.58 | 1.99E-02 |
| | | | hsa-miR-12135;iso_snp:8CT;iso_5p:t | -1.66 | 8.20E-02 |
| | | | hsa-miR-4448;iso_snp:6GC | -2.17 | 1.67E-04 |

| LEV | | | | | |
|---|---|---|---|---|---|
| Upregulated miRNA | | | Downregulated miRNAs | | |
| Name | Log2Change | padj | Name | Log2Change | padj |
| hsa-miR-495-3p | 3.77 | 2.01E-02 | hsa-miR-425-5p;iso_5p:a | -0.60 | 9.90E-02 |
| hsa-miR-136-3p | 2.54 | 8.30E-02 | hsa-miR-23b-3p | -0.84 | 9.90E-02 |
| hsa-miR-376a-3p | 2.18 | 8.36E-02 | hsa-miR-10a-5p;iso_5p:t | -1.45 | 8.30E-02 |
| hsa-miR-136-5p | 2.13 | 2.01E-02 | hsa-miR-192-5p;iso_5p:c | -1.47 | 9.90E-02 |
| hsa-miR-376c-3p | 2.06 | 4.25E-02 | hsa-miR-378a-3p | -1.55 | 1.32E-02 |
| hsa-miR-369-3p | 1.95 | 5.90E-02 | hsa-miR-502-3p | -1.59 | 8.30E-02 |
| hsa-miR-424-5p | 1.67 | 3.66E-02 | hsa-miR-199a-5p | -1.80 | 1.32E-02 |
| hsa-miR-493-5p | 1.65 | 8.30E-02 | hsa-miR-214-3p | -1.96 | 7.67E-02 |
| hsa-miR-27a-3p | 1.08 | 2.48E-02 | hsa-miR-10b-5p | -2.61 | 5.90E-02 |
| | | | hsa-miR-12136 | -3.20 | 7.28E-05 |

| SEV | | | | | |
|---|---|---|---|---|---|
| Upregulated miRNAs | | | Downregulated miRNAs | | |
| Name | Log2Change | padj | Name | Log2Change | padj |
| hsa-miR-665 | 5.02 | 2.75E-04 | hsa-miR-425-5p;iso_5p:a | -0.45 | 1.65E-02 |
| hsa-miR-655-3p | 2.83 | 1.26E-02 | hsa-miR-23b-3p | -0.65 | 1.28E-02 |
| hsa-miR-4791 | 2.71 | 1.90E-02 | hsa-miR-30c-5p | -0.78 | 7.78E-03 |
| hsa-miR-323b-3p | 2.26 | 4.36E-03 | hsa-miR-10a-5p | -0.95 | 2.31E-02 |
| hsa-miR-154-5p | 1.86 | 1.06E-03 | hsa-miR-10a-3p | -1.16 | 4.21E-02 |
| hsa-miR-377-3p | 1.74 | 1.15E-02 | hsa-miR-10a-5p;iso_5p:t | -1.23 | 4.34E-03 |
| hsa-miR-495-3p | 1.69 | 1.19E-02 | hsa-miR-12136 | -1.28 | 2.02E-02 |
| hsa-miR-889-3p | 1.64 | 1.59E-02 | hsa-miR-7977;iso_snp:5GA;iso_5p:t | -1.60 | 1.37E-02 |
| hsa-miR-376a-3p | 1.59 | 6.08E-03 | hsa-miR-3200-3p | -2.44 | 1.65E-02 |
| hsa-miR-376a-3p;iso_snp:6[illegible] | 1.27 | 1.31E-02 | hsa-miR-4516;iso_snp:16GA;iso_5p:gg | -2.82 | 9.75E-03 |
| | | | hsa-miR-4516;iso_snp:15GA;iso_5p:ggg | -3.15 | 6.22E-03 |

Table S1: miRNAs differentially expressed (upregulated in green, downregulated in orange) between healthy and COPD SA-epithelial cells.

| SAF | | | | | |
|---|---|---|---|---|---|
| **Upregulated miRNAs** | | | **Downregulated miRNAs** | | |
| **Name** | **Log2Change** | **padj** | **Name** | **Log2Change** | **padj** |
| hsa-miR-210-3p | 2.63 | 6.26E-07 | hsa-miR-425-5p | -0.39 | 3.41E-02 |
| hsa-miR-3178;iso_snp:1AG;iso_5p:GC | 1.92 | 1.71E-02 | hsa-miR-576-5p | -0.49 | 3.84E-02 |
| hsa-miR-188-5p | 1.83 | 1.56E-03 | hsa-miR-151a-3p | -0.54 | 4.78E-02 |
| hsa-miR-204-5p | 1.82 | 3.37E-03 | hsa-miR-197-3p | -0.64 | 1.68E-02 |
| hsa-miR-1-3p | 1.58 | 4.76E-04 | hsa-miR-17-5p | -0.65 | 1.93E-02 |
| hsa-miR-490-3p | 1.58 | 7.00E-03 | hsa-let-7c-5p | -0.72 | 1.81E-02 |
| hsa-miR-450a-2-3p;iso_5p:a | 1.48 | 1.22E-02 | hsa-miR-99a-5p | -0.83 | 2.32E-02 |
| hsa-miR-376a-5p | 1.44 | 4.37E-03 | hsa-miR-625-3p | -0.84 | 4.72E-02 |
| hsa-miR-887-3p | 1.39 | 4.42E-02 | hsa-miR-192-5p;iso_5p:c | -0.92 | 2.79E-02 |
| hsa-miR-369-3p | 1.34 | 7.61E-03 | hsa-miR-155-5p | -1.16 | 1.67E-02 |
| hsa-miR-1277-3p | 1.31 | 2.83E-02 | hsa-miR-941 | -1.33 | 2.39E-03 |
| hsa-miR-127-5p | 1.29 | 5.58E-03 | hsa-miR-3928-3p | -1.61 | 3.45E-02 |
| hsa-miR-98-3p | 1.19 | 3.30E-02 | hsa-miR-7704;iso_5p:c | -5.36 | 3.66E-04 |
| hsa-miR-136-5p | 1.15 | 6.59E-03 | | | |
| hsa-let-7i-3p | 1.11 | 7.92E-05 | | | |
| hsa-miR-7-1-3p | 1.08 | 1.16E-02 | | | |
| hsa-miR-193a-3p | 1.06 | 4.67E-02 | | | |
| hsa-miR-137-3p | 1.05 | 6.16E-03 | | | |
| hsa-miR-143-5p | 1.02 | 4.36E-03 | | | |
| hsa-miR-376b-3p | 1.02 | 2.57E-02 | | | |
| hsa-miR-1307-5p | 0.99 | 1.68E-02 | | | |
| hsa-miR-299-5p | 0.99 | 1.66E-02 | | | |
| hsa-miR-136-3p;iso_5p:c | 0.91 | 4.14E-02 | | | |
| hsa-miR-889-3p | 0.84 | 4.12E-02 | | | |
| hsa-miR-450a-1-3p | 0.81 | 2.64E-02 | | | |
| hsa-miR-29c-5p | 0.81 | 1.40E-02 | | | |
| hsa-miR-152-3p | 0.79 | 5.41E-03 | | | |
| hsa-miR-22-3p | 0.78 | 2.58E-03 | | | |
| hsa-miR-22-5p | 0.76 | 2.49E-03 | | | |
| hsa-miR-127-5p;iso_5p:ct | 0.74 | 3.79E-02 | | | |
| hsa-miR-21-3p | 0.70 | 1.83E-02 | | | |
| hsa-miR-497-5p | 0.68 | 1.68E-02 | | | |
| hsa-miR-1185-1-3p | 0.66 | 4.60E-02 | | | |
| hsa-miR-21-5p | 0.65 | 2.51E-02 | | | |
| hsa-miR-186-5p | 0.58 | 1.21E-02 | | | |
| hsa-miR-34a-5p | 0.55 | 2.70E-02 | | | |
| hsa-miR-27a-3p | 0.53 | 4.72E-02 | | | |
| hsa-miR-628-5p | 0.53 | 3.11E-02 | | | |
| hsa-miR-148b-3p | 0.48 | 4.71E-02 | | | |

| LEVs | | | | | |
|---|---|---|---|---|---|
| Upregulated miRNAs | | | Downregulated miRNAs | | |
| Name | Log2Change | padj | Name | Log2Change | padj |
| hsa-miR-1185-2-3p | 3.16 | 4.57E-02 | hsa-miR-128-3p | -0.50 | 9.21E-03 |
| hsa-miR-210-3p | 2.08 | 1.08E-04 | hsa-miR-629-5p | -0.63 | 3.18E-02 |
| hsa-miR-544a | 2.05 | 9.03E-03 | hsa-miR-29a-3p | -0.67 | 2.85E-02 |
| hsa-miR-27b-5p | 1.72 | 6.69E-03 | hsa-miR-99a-5p | -0.84 | 2.92E-02 |
| hsa-miR-204-5p | 1.59 | 2.10E-02 | hsa-miR-378a-3p | -0.88 | 1.97E-02 |
| hsa-miR-433-3p | 1.55 | 7.93E-03 | hsa-miR-17-3p | -0.93 | 4.68E-02 |
| hsa-miR-452-3p;iso_5p:TCAGT | 1.55 | 3.81E-02 | | | |
| hsa-miR-376a-5p | 1.45 | 1.78E-02 | | | |
| hsa-miR-193a-5p | 1.34 | 3.40E-02 | | | |
| hsa-miR-29b-2-5p | 1.29 | 3.32E-02 | | | |
| hsa-miR-299-5p | 1.21 | 7.31E-03 | | | |
| hsa-miR-376b-3p | 1.10 | 3.39E-02 | | | |
| hsa-miR-137-3p | 0.88 | 3.15E-02 | | | |
| hsa-let-7d-3p | 0.84 | 7.97E-03 | | | |

| SEVs | | | | | |
|---|---|---|---|---|---|
| Upregulated miRNAs | | | Downregulated miRNAs | | |
| Name | Log2Change | padj | Name | Log2Change | padj |
| hsa-miR-137-5p | 3.09 | 8.43E-03 | hsa-miR-16-5p | -0.58 | 4.29E-02 |
| hsa-miR-154-3p | 1.91 | 1.14E-02 | hsa-miR-1255b-5p | -4.20 | 5.34E-04 |
| hsa-miR-296-5p;iso_5p:G | 1.81 | 2.75E-02 | hsa-miR-93-5p | -0.67 | 1.92E-02 |
| hsa-miR-410-3p | 1.79 | 2.98E-02 | hsa-miR-151a-3p | -0.73 | 8.65E-03 |
| hsa-miR-27b-5p | 1.71 | 1.38E-02 | hsa-miR-629-5p | -0.73 | 9.08E-03 |
| hsa-miR-376a-5p | 1.55 | 1.79E-02 | hsa-miR-505-3p;iso_5p:c | -0.99 | 2.72E-02 |
| hsa-miR-204-5p | 1.52 | 2.63E-02 | hsa-miR-1246 | -1.71 | 1.53E-02 |
| hsa-miR-433-3p | 1.44 | 1.39E-02 | hsa-miR-3928-3p | -2.43 | 2.25E-02 |
| hsa-let-7f-1-3p | 1.11 | 3.62E-02 | hsa-miR-665 | -2.64 | 1.11E-02 |
| hsa-miR-487b-3p | 1.08 | 9.49E-03 | hsa-miR-1299 | -3.09 | 6.86E-03 |
| hsa-miR-323a-3p;iso_5p:G | 1.02 | 2.35E-02 | hsa-miR-5699-3p;iso_5p:t | -3.31 | 1.60E-03 |
| hsa-miR-1185-1-3p | 0.98 | 1.84E-02 | | | |
| hsa-miR-137-3p | 0.97 | 1.24E-02 | | | |
| hsa-miR-369-5p | 0.88 | 3.14E-02 | | | |
| hsa-miR-154-5p | 0.85 | 3.63E-02 | | | |
| hsa-miR-145-5p | 0.83 | 4.84E-02 | | | |
| hsa-let-7e-3p | 0.81 | 4.79E-02 | | | |
| hsa-miR-487b-3p;iso_5p:aa | 0.78 | 4.82E-02 | | | |
| hsa-miR-374b-5p | 0.64 | 4.06E-02 | | | |
| hsa-miR-22-3p | 0.59 | 2.24E-02 | | | |

Table S2: miRNAs differentially expressed (upregulated in green, downregulated in orange) between healthy and COPD SA-fibroblasts

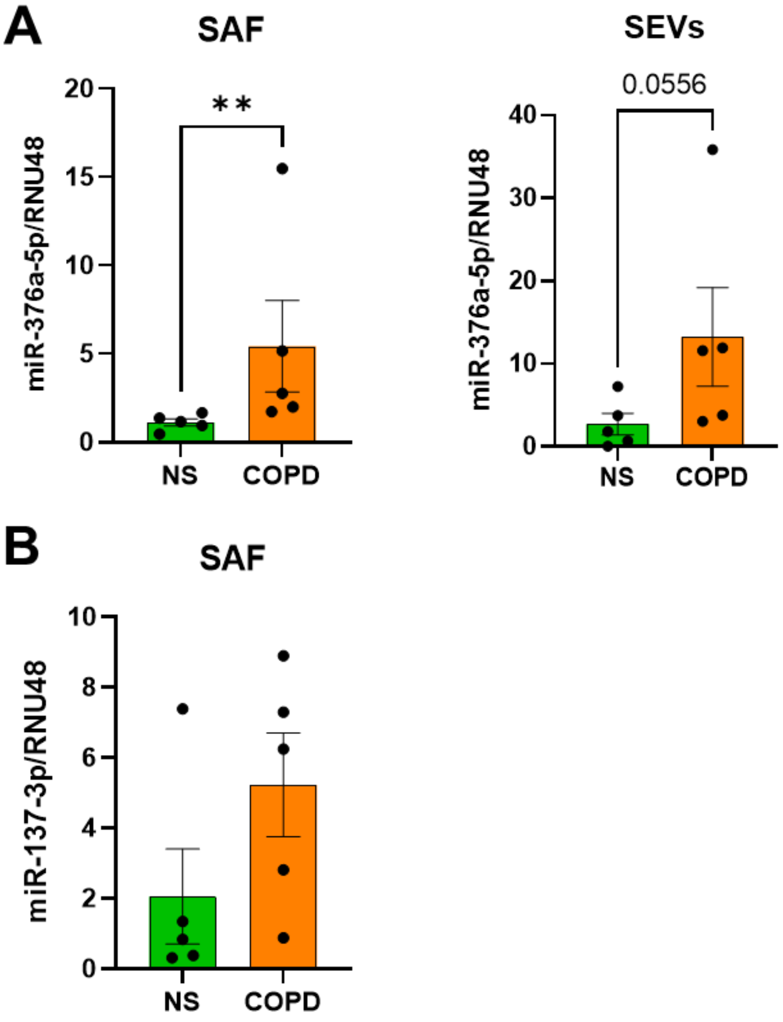


<u>Figure S1: miR-376a-5p and miR-137-3p are upregulated in COPD SA-fibroblasts</u>

Total RNA was isolated from healthy and COPD SA-fibroblasts and the expression of **A** miR-376a-5p and **B** miR-137-3p was analysed by RT-qPCR (n=5). Data are represented as mean ± SEM and analysed by Mann-Whitney test. **p<0.01

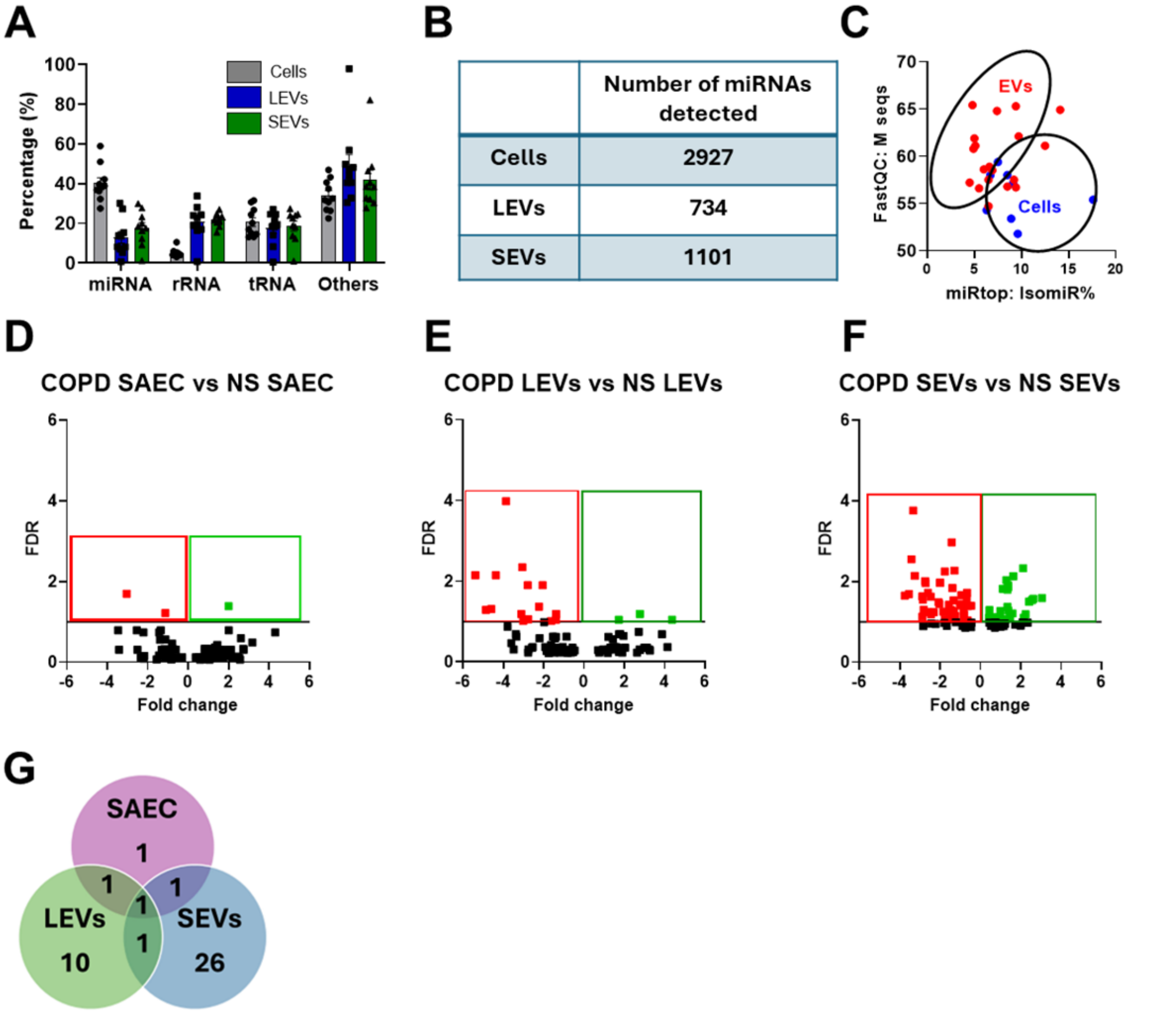


<u>Figure S2: Dysregulation of miRNA Expression in COPD SA-epithelial cells and Associated EVs in response to oxidative stress.</u>

Small RNA sequencing was conducted on healthy, and COPD SA-epithelial cells cultured with 100µM of $H_2O_2$ and their derived large and small EVs. **A.** The proportions of miRNA, rRNA, tRNA and other types of smRNA were characterised and **B.** the numbers of miRNAs detected in both SAEC and EVs were analysed. **C.** The percentage of IsomiR (variants of miRNAs) and Mseq (sequence count) were plotted to examine the miRNAs profiles in cells and EVs. **D-F.** Volcano plots were used to compare miRNA expression between healthy and COPD cells and EVs associated. **G.** A Venn-diagram illustrated the overlap of dysregulation miRNA between healthy and COPD cells and EVs. **H.** The sequencing data were validated by RT-qPCR for miR-376a-3p (n=5). Padj value<0.1. Data are represented as mean ± SEM and analysed by Mann-Whitney test. *p<0.05

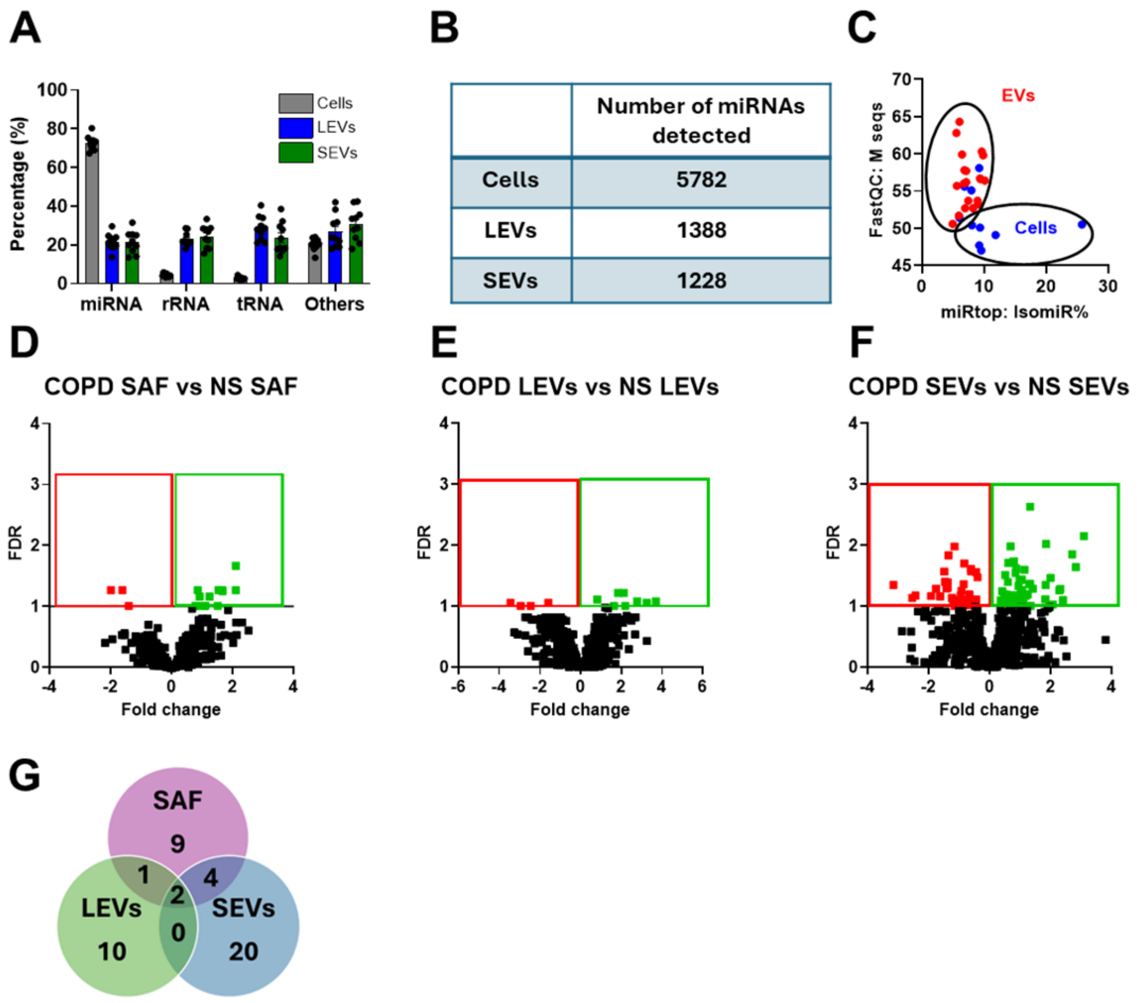


<u>Figure S3: Dysregulation of miRNA Expression in COPD SA-fibroblasts and Associated EVs in response to oxidative stress.</u>

Small RNA sequencing was conducted on healthy and COPD SA-fibroblasts cultured with 100µM of $H_2O_2$ and their derived large and small EVs. **A.** The proportions of miRNA, rRNA, tRNA and other types of smRNA were characterised and **B.** the numbers of miRNAs detected in both SAF and EVs were analysed. **C.** The percentage of IsomiR (variants of miRNAs) and Mseq (sequence count) were plotted to examine the miRNAs profiles in cells and EVs. **D-F.** Volcano plots were used to compare miRNA expression between healthy and COPD cells and EVs associated. **G.** A Venn-diagram illustrated the overlap of dysregulation miRNA between healthy and COPD cells and EVs. **H.** The sequencing data were validated by RT-qPCR for miR-376a-3p (n=5). Padj value<0.1. Data are represented as mean ± SEM and analysed by Mann-Whitney test. *p<0.05

| Healthy SAEC | | | | | |
|---|---|---|---|---|---|
| Upregulated in response to $H_2O_2$ | | | Downregulated in response to $H_2O_2$ | | |
| | Log2Change | pvalue | Name | Log2Change | pvalue |
| hsa-miR-194-5p | 0.71 | 3.03E-02 | hsa-miR-339-5p | -0.50 | 7.71E-02 |
| hsa-miR-142-3p | 0.67 | 9.19E-02 | hsa-miR-4448;iso_snp:6GC | -0.72 | 5.33E-02 |
| hsa-miR-424-5p | 0.67 | 9.14E-02 | hsa-miR-3178;iso_snp:1AG;iso_5p:GC | -1.08 | 7.20E-02 |
| hsa-miR-96-5p | 0.62 | 5.27E-02 | hsa-miR-4454;iso_5p:ggat | -1.56 | 2.89E-02 |
| hsa-miR-27a-3p | 0.60 | 6.76E-02 | hsa-miR-7704;iso_5p:c | -2.03 | 6.38E-03 |
| hsa-miR-30b-5p | 0.59 | 9.29E-02 | hsa-miR-92b-5p | -2.69 | 2.90E-02 |

| Healthy LEVs | | | | | |
|---|---|---|---|---|---|
| Upregulated in response to $H_2O_2$ | | | Downregulated in response to $H_2O_2$ | | |
| | Log2Change | pvalue | Name | Log2Change | pvalue |
| hsa-miR-148b-5p;iso_5p:G | 1.05 | 9.27E-02 | hsa-miR-181c-3p;iso_5p:a | -4.28 | 4.86E-03 |
| hsa-miR-652-3p | 1.26 | 1.63E-02 | hsa-let-7f-1-3p | -3.26 | 3.70E-02 |
| hsa-miR-324-3p;iso_5p:c | 1.50 | 4.25E-02 | hsa-let-7i-3p | -3.19 | 3.26E-02 |
| hsa-miR-501-5p | 1.93 | 9.86E-02 | hsa-miR-590-3p;iso_5p:G | -3.10 | 5.35E-02 |
| hsa-miR-23a-5p | 2.45 | 6.45E-02 | hsa-miR-455-5p;iso_5p:ta | -1.02 | 6.05E-02 |
| hsa-miR-4791 | 2.64 | 2.10E-02 | | | |
| hsa-miR-708-3p | 2.90 | 2.67E-02 | | | |

| Healthy SEVs | | | | | |
|---|---|---|---|---|---|
| Upregulated in response to $H_2O_2$ | | | Downregulated in response to $H_2O_2$ | | |
| | Log2Change | pvalue | Name | Log2Change | pvalue |
| hsa-miR-324-5p | 0.764566652 | 0.038769543 | hsa-miR-7976 | -2.622438602 | 0.067775 |
| hsa-miR-941 | 1.141586302 | 0.020709323 | hsa-miR-3200-3p | -2.431324535 | 0.034156 |
| hsa-miR-339-3p | 1.146516534 | 0.017215801 | hsa-miR-139-5p | -1.823865891 | 0.04547 |
| hsa-miR-548j-5p | 1.491818055 | 0.07778144 | hsa-miR-151a-5p | -0.546379995 | 0.088338 |
| hsa-miR-1285-3p | 3.081990612 | 0.064691647 | hsa-miR-126-3p | -0.491450573 | 0.091793 |

| COPD SAEC | | | | | |
|---|---|---|---|---|---|
| Upregulated in response to $H_2O_2$ | | | Downregulated in response to $H_2O_2$ | | |
| | Log2Change | pvalue | Name | Log2Change | pvalue |
| hsa-miR-194-5p | 0.40 | 8.36E-02 | hsa-miR-17-3p | -0.61 | 5.48E-02 |
| hsa-let-7d-3p | 0.59 | 4.83E-02 | hsa-miR-361-5p | -0.34 | 7.84E-02 |
| hsa-miR-10401-3p;iso_5p:G | 0.63 | 5.70E-02 | hsa-miR-29a-3p | -0.31 | 7.25E-02 |
| hsa-miR-142-3p | 0.73 | 6.27E-02 | hsa-miR-103a-3p | -0.28 | 9.92E-02 |
| hsa-miR-12135;iso_snp:8CT;iso_5p:t | 0.77 | 3.44E-03 | | | |
| hsa-miR-154-3p | 1.33 | 8.64E-02 | | | |

| COPD LEVs | | | | | |
|---|---|---|---|---|---|
| Upregulated in response to $H_2O_2$ | | | Downregulated in response to $H_2O_2$ | | |
| | Log2Change | pvalue | Name | Log2Change | pvalue |
| hsa-miR-1246;iso_5p:a | 0.40 | 7.03E-02 | hsa-miR-337-3p;iso_5p:c | -3.54 | 5.67E-02 |
| hsa-miR-16-5p | 0.44 | 5.05E-02 | hsa-miR-4497;iso_5p:c | -1.60 | 4.71E-04 |
| hsa-miR-190b-5p | 1.28 | 7.14E-02 | hsa-miR-145-5p | -0.90 | 1.86E-02 |
| hsa-miR-1-3p | 1.52 | 1.24E-02 | hsa-miR-3178;iso_snp:1AG;iso_5p:GC | -0.76 | 3.74E-02 |
| hsa-miR-4791 | 1.54 | 8.50E-02 | hsa-miR-17-5p | -0.59 | 5.82E-02 |
| hsa-miR-502-3p | 1.88 | 9.71E-02 | hsa-miR-29b-3p | -0.47 | 5.20E-02 |

| COPD SEVs | | | | | |
|---|---|---|---|---|---|
| Upregulated in response to $H_2O_2$ | | | Downregulated in response to $H_2O_2$ | | |
| **Name** | **Log2Change** | **pvalue** | **Name** | **Log2Change** | **pvalue** |
| hsa-miR-29b-3p | 0.32 | 8.30E-03 | hsa-miR-3940-5p;iso_snp:8CT;iso_5p:G | -1.59 | 7.00E-02 |
| hsa-miR-424-5p | 0.47 | 7.40E-03 | hsa-miR-203a-3p | -0.43 | 1.00E-02 |
| hsa-miR-369-3p | 0.48 | 3.85E-02 | | | |
| hsa-miR-340-5p | 0.50 | 9.75E-02 | | | |
| hsa-miR-10401-3p;iso_5p:G | 0.69 | 9.66E-02 | | | |
| hsa-miR-153-3p | 1.31 | 8.01E-03 | | | |
| hsa-miR-296-5p | 2.01 | 4.37E-02 | | | |

Table S3: miRNAs differentially expressed (upregulated in green, downregulated in orange) between baseline and $H_2O_2$ conditions in healthy and COPD SA-epithelial cells.

| Healthy SAF | | | | | |
|---|---|---|---|---|---|
| Upregulated in response to $H_2O_2$ | | | Downregulated in response to $H_2O_2$ | | |
| Name | Log2Change | pvalue | Name | Log2Change | pvalue |
| hsa-miR-3178;iso_snp:1AG;iso_5p:GC | -1.90282566 | 4.40E-02 | hsa-miR-155-5p | 0.26 | 2.20E-02 |
| hsa-miR-769-3p | -1.518929 | 6.66E-02 | hsa-miR-17-5p | 0.32 | 8.23E-02 |
| hsa-let-7a-3p | -0.39407343 | 4.85E-02 | hsa-miR-503-3p;iso_5p:g | 1.49 | 5.24E-02 |

| Healthy LEVs | | | | | |
|---|---|---|---|---|---|
| Upregulated in response to $H_2O_2$ | | | Downregulated in response to $H_2O_2$ | | |
| Name | Log2Change | pvalue | Name | Log2Change | pvalue |
| hsa-miR-29b-1-5p | 0.92 | 2.63E-02 | hsa-miR-545-3p | -2.45 | 8.80E-02 |
| hsa-miR-4497;iso_5p:ct | 1.32 | 9.61E-02 | hsa-miR-95-3p | -2.40 | 4.27E-02 |
| hsa-miR-27a-5p | 1.51 | 4.13E-02 | hsa-miR-103a-2-5p | -2.13 | 8.70E-02 |
| hsa-miR-3605-3p | 1.73 | 2.03E-02 | hsa-miR-664b-3p | -1.93 | 9.17E-02 |
| hsa-miR-30c-1-3p | 2.07 | 5.48E-02 | hsa-miR-452-5p;iso_5p:aac | -1.09 | 9.08E-02 |
| hsa-miR-1185-2-3p | 3.57 | 1.74E-02 | hsa-miR-136-3p | -1.03 | 6.15E-02 |
| | | | hsa-miR-31-3p | -0.96 | 3.65E-02 |
| | | | hsa-let-7a-3p | -0.73 | 6.66E-02 |

| Healthy SEVs | | | | | |
|---|---|---|---|---|---|
| Upregulated in response to $H_2O_2$ | | | Downregulated in response to $H_2O_2$ | | |
| Name | Log2Change | pvalue | Name | Log2Change | pvalue |
| hsa-miR-203a-3p | 0.64 | 8.79E-02 | hsa-miR-1255b-5p | -2.88 | 7.07E-03 |
| hsa-miR-548aq-3p | 1.95 | 7.32E-02 | hsa-miR-1246;iso_5p:a | -0.91 | 5.79E-03 |
| | | | hsa-miR-183-5p | -0.68 | 9.10E-02 |
| | | | hsa-miR-1246;iso_5p:aa | -0.64 | 3.85E-02 |

| COPD SAF | | | | | |
|---|---|---|---|---|---|
| Upregulated in response to $H_2O_2$ | | | Downregulated in response to $H_2O_2$ | | |
| | Log2Change | pvalue | Name | Log2Change | pvalue |
| hsa-miR-192-5p | 0.40 | 6.69E-02 | hsa-miR-130a-5p | -2.90 | 3.87E-02 |
| | | | hsa-miR-496;iso_5p:tg | -1.35 | 6.87E-02 |
| | | | hsa-miR-941 | -0.36 | 6.44E-02 |

| COPD LEVs | | | | | |
|---|---|---|---|---|---|
| Upregulated in response to $H_2O_2$ | | | Downregulated in response to $H_2O_2$ | | |
| | Log2Change | pvalue | Name | Log2Change | pvalue |
| hsa-miR-1307-5p | 0.46 | 9.07E-02 | hsa-miR-10401-3p;iso_5p:G | -0.97 | 6.89E-02 |
| hsa-miR-27a-5p | 1.32 | 1.58E-02 | | | |
| hsa-miR-99a-3p | 1.65 | 3.42E-02 | | | |

| COPD SEVs | | | | | |
|---|---|---|---|---|---|
| Upregulated in response to $H_2O_2$ | | | Downregulated in response to $H_2O_2$ | | |
| **Name** | **Log2Change** | **pvalue** | **Name** | **Log2Change** | **pvalue** |
| hsa-miR-16-5p | 0.32 | 3.74E-02 | hsa-miR-4497;iso_5p:ct | -1.10 | 9.90E-03 |
| hsa-miR-29b-3p | 0.32 | 5.08E-02 | hsa-miR-203a-3p | -0.89 | 4.16E-02 |
| hsa-miR-26b-5p | 0.34 | 3.60E-02 | hsa-miR-486-5p | -0.65 | 1.23E-02 |
| hsa-miR-20a-5p | 0.36 | 2.97E-02 | hsa-miR-485-3p | -0.62 | 6.44E-02 |
| hsa-miR-30e-5p | 0.42 | 9.87E-03 | hsa-miR-4454;iso_5p:ggat | -0.58 | 6.97E-02 |
| hsa-miR-15b-5p | 0.46 | 2.90E-02 | hsa-miR-323b-3p | -0.55 | 9.46E-02 |
| hsa-miR-19b-3p | 0.50 | 2.01E-02 | hsa-miR-125b-5p | -0.50 | 4.01E-03 |
| hsa-miR-454-3p | 0.60 | 2.35E-03 | hsa-miR-134-5p | -0.43 | 3.99E-02 |
| hsa-miR-301a-3p | 0.61 | 4.50E-02 | hsa-miR-432-5p | -0.43 | 3.99E-02 |
| hsa-miR-3613-5p | 0.69 | 4.46E-02 | hsa-miR-431-5p | -0.42 | 7.66E-02 |
| hsa-miR-200a-3p | 0.72 | 1.27E-02 | hsa-miR-193a-5p | -0.42 | 5.42E-02 |
| hsa-miR-340-5p | 0.75 | 1.89E-02 | hsa-miR-127-3p | -0.41 | 1.33E-02 |
| hsa-miR-15b-3p | 0.80 | 2.89E-02 | hsa-miR-409-3p | -0.39 | 5.83E-02 |
| hsa-miR-192-5p | 0.94 | 1.32E-02 | hsa-miR-342-3p | -0.36 | 2.48E-02 |
| hsa-miR-452-5p;iso_5p:aac | 1.05 | 4.55E-03 | hsa-miR-92a-3p | -0.36 | 5.20E-02 |
| | | | hsa-miR-222-3p | -0.35 | 4.41E-02 |
| | | | hsa-miR-320a-3p | -0.33 | 7.21E-02 |
| | | | hsa-miR-99b-5p | -0.27 | 9.07E-02 |

Table S4: miRNAs differentially expressed (upregulated in green, downregulated in orange) between baseline and $H_2O_2$ conditions in healthy and COPD SA-fibroblasts.
